## Supplemental Information for "Development and Assessment of a Sustainable PhD Internship Program Supporting Diverse Biomedical Career Outcomes"

**Logistic Regression Posthoc Model With Social Identity Groups Included (Model 2)**

Post hoc tests explored the potential impact of social identity groups on career matching, but trends were nearly identical, hence the original, simple Model 1 was retained. Model 2 included demographic information [Race/Ethnicity (UR/WR), Gender (Female/Male), and Citizenship (Citizen/International)]. Race/Ethnicity and Citizenship did not impact the model, indicating that UR and international applicants were as likely as WR and citizen applicants to match their career interests (p<0.52, OR=1.00 and p<0.26, OR=1.00, respectively). For this reason, we ran a simplified model including only Gender as a demographic variable. The trends were similar for Career interests and Trainee Type (Supplemental Table 1). The overall model remained statistically significant while controlling for Gender in addition to the trainee type, and career interests, with nearly identical patterns and significance levels for the impact participation (OR=3.50, p<0.001) (χ^2^ = 97.57, R^2^_Nagelkerke_ = 0.21, p<0.001). Interestingly, applicants who identified as Female were nearly one and a half times more likely to match their career interests when compared with applicants who identified as Male (p=0.02, OR=1.57).

**Supplemental Table 1.** **Summary for regression model controlling for demographic variables Race/Ethnicity (UR/WR), Gender (Female/Male), and Citizenship (Citizen/International)**

| **Variables** | p-value | Exp(B) | 95% C.I. for EXP(B) | |
| --- | --- | --- | --- | --- |
|  |  |  | Lower | Upper |
| **Internship Participation***** | **p<0.001** | **3.51** | **1.71** | **7.21** |
| **Number of Career Interests***** | **p<0.001** | **1.20** | **1.11** | **1.29** |
| **Trainee Type (Pd)***** | **p<0.001** | **5.03** | **2.94** | **8.60** |
| **Gender*** | **p=0.02** | **1.58** | **1.07** | **2.33** |
| Race/Ethnicity (ns) | p=0.52 | 1.00 | 1.00 | 1.01 |
| Citizenship (ns) | p=0.26 | 1.00 | 0.99 | 1.00 |
